# Spatial and temporal context jointly modulate the sensory response within the ventral visual stream

**DOI:** 10.1101/2020.07.24.219709

**Authors:** Tao He, David Richter, Zhiguo Wang, Floris P. de Lange

**Affiliations:** School of Psychological and Cognitive Sciences and Beijing Key Laboratory of Behavior and Mental Health, Peking University, Beijing 100871, China; IDG/McGovern Institute for Brain Research, Peking University, Beijing 100871, China; Donders Institute for Brain, Cognition and Behaviour, Radboud University, Kapittelweg 29, 6525 EN Nijmegen, the Netherlands; Institutes of Psychological Sciences, Hangzhou Normal University, Hangzhou, 311121, China

## Abstract

Both spatial and temporal context play an important role in visual perception and behavior. Humans can extract statistical regularities from both forms of context to help processing the present and to construct expectations about the future. Numerous studies have found reduced neural responses to expected stimuli compared to unexpected stimuli, for both spatial and temporal regularities. However, it is largely unclear whether and how these forms of context interact. In the current fMRI study, thirty-three human volunteers were exposed to object stimuli that could be expected or surprising in terms of their spatial and temporal context. We found a reliable independent contribution of both spatial and temporal context in modulating the neural response. Specifically, neural responses to stimuli in expected compared to unexpected contexts were suppressed throughout the ventral visual stream. Interestingly, the modulation by spatial context was stronger in magnitude and more reliable than modulations by temporal context. These results suggest that while both spatial and temporal context serve as a prior that can modulate sensory processing in a similar fashion, predictions of spatial context may be a more powerful modulator in the visual system.

**Significance Statement:** Both temporal and spatial context can affect visual perception, however it is largely unclear if and how these different forms of context interact in modulating sensory processing. When manipulating both temporal and spatial context expectations, we found that they jointly affected sensory processing, evident as a suppression of neural responses for expected compared to unexpected stimuli. Interestingly, the modulation by spatial context was stronger than that by temporal context. Together, our results suggest that spatial context may be a stronger modulator of neural responses than temporal context within the visual system. Thereby, the present study provides new evidence how different types of predictions jointly modulate perceptual processing.

## Introduction

Humans are exquisitely sensitive to visual statistical regularities. Indeed, knowledge of both spatial and temporal context can facilitate visual perception and perceptual decision-making (Bar, 2004). For instance, in the case of spatial context, a foreground object is more easily identified when it appears on congruent backgrounds, compared to when it appears on incongruent backgrounds (Davenport and Potter, 2004). Facilitatory effects of temporal context have also been shown, for instance during exposure to sequentially presented stimuli, with faster and more accurate responses to expected compared to unexpected stimuli (Bertels et al., 2012; Hunt & Aslin, 2001; Richter & de Lange, 2019). At the same time neural responses have been shown to be modulated by temporal context, with a marked suppression of sensory responses to expected compared to unexpected stimuli, reported in humans (Summerfield et al., 2008; den Ouden et al., 2009; Egner et al., 2010; Richter et al., 2018; Richter and de Lange, 2019) and non-human primates (Freedman et al., 2006; Meyer and Olson, 2011; Kaposvari et al., 2018). However, comparatively less is known about the modulation of neural responses by spatial context. Human fMRI studies suggest that a similar network of (sub-)cortical areas is involved in learning spatial contexts as during learning of temporal sequences (Karuza et al., 2017). Thus, while the learning process of temporal and spatial regularities may share neural characteristics, the consequences for sensory processing, following the acquisition of spatial regularities, remain unknown. In particular, do predictions of spatial context result in a similar suppression of neural responses as temporal sequence predictions? Moreover, it is currently unclear if and how spatial and temporal context may interact in sharping sensory processing.

In the current study, we set out to concurrently examine the neural and behavioral consequences of spatial and temporal contextual expectations following statistical learning. To this end, participants were exposed to leading image pairs, consisting of two object images presented left and right of fixation, which predicted the identity of trailing object image pairs, thus rendering the trailing images expected based on the temporal context. Moreover, the simultaneously presented images were also predictive of each other, thus generating a predictable spatial context (see **Figure 1c**). Blood oxygenation level-dependent (BOLD) signals were recorded with functional magnetic resonance imaging (fMRI), while participants monitored the images for occasional target images (i.e., flipped object images) that occurred at unpredictable moments.

**Figure 1.**
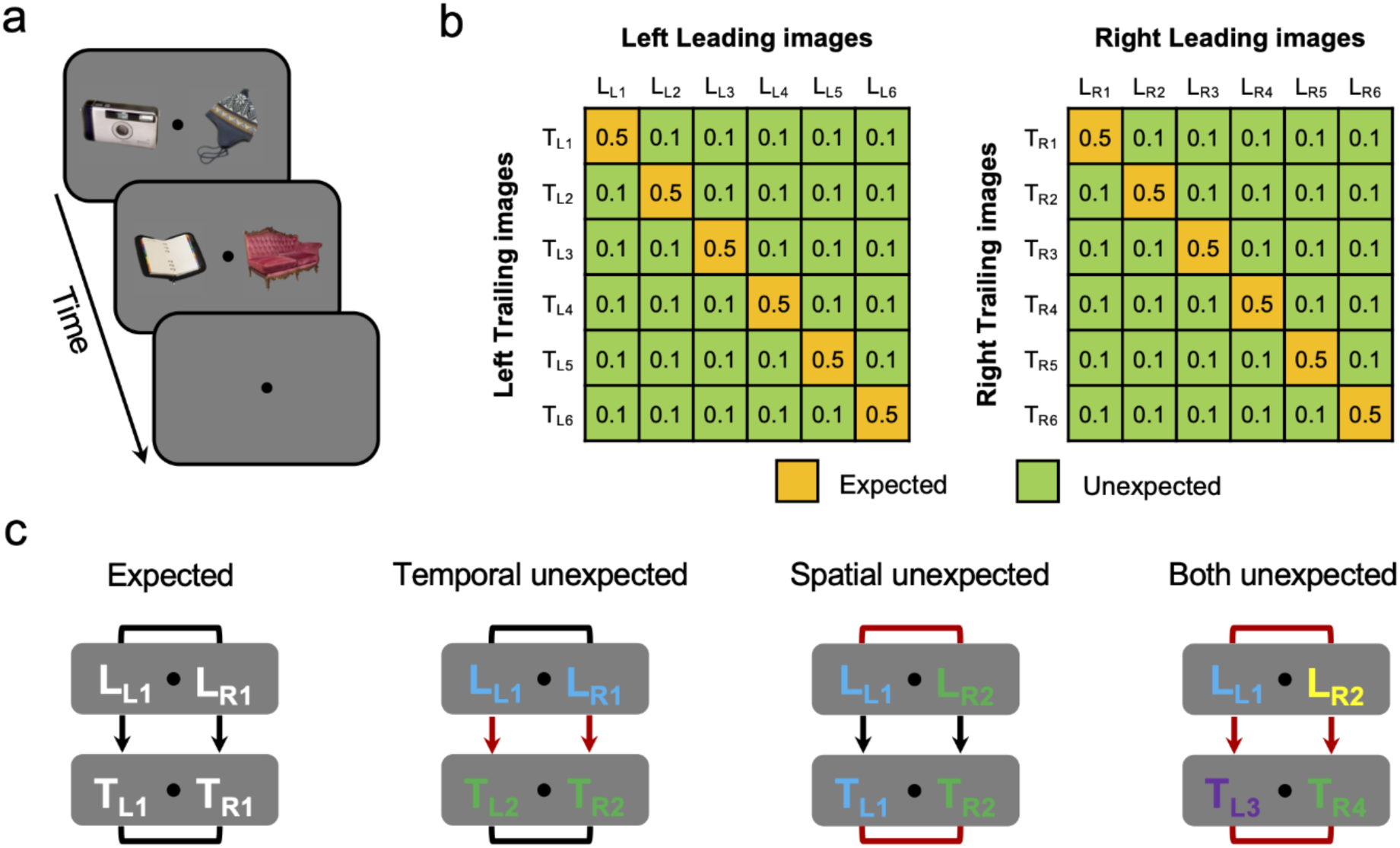
Experimental paradigm and design. **(a)** Experimental paradigm in both the behavioral learning and fMRI session. A trial starts with a 500 ms presentation of two leading images, presented left and right from the central fixation dot. The two leading images are immediately followed by the trailing images, without ISI, at the same locations, also shown for 500 ms. Participants were asked to detect an infrequently presented upside-down version of the images (∼10% of trials). Trials were separated by a 2 - 6 s (mean 3 s) ITI period. **(b)** Shown are the image transition matrices determining the statistical regularities between leading and trailing images during MRI scanning. On the left, L_L1_ to L_L6_ represent the six leading images presented on the left of the fixation dot, while T_L1_ to T_L6_ represent the associated six left trailing images. Similarly, L_R1_ to L_R6_ represent the six right leading images, while T_R1_ to T_R6_ represent the six right trailing images. Yellow cells indicate image pairs that are expected by temporal context, while green denotes unexpected image pairs. Numbers represent the probability of that cell during MRI scanning. Crucially, the left and right images were also associated with each other, constituting the spatial context. For instance, L_L1_ was associated with L_R1_, and T_L1_ was associated with T_R1_. In this case, L_L1_, L_R1_, T_L1_ and T_R1_ composed two image pairs that were expected in both the temporal and spatial contexts (see **Figure1c**, ‘Expected’). **(c)** Illustration of the four expectation conditions during MRI scanning. Black lines indicate expected associations, while red lines indicate unexpected pairings. *Expected condition*: the matched image configuration that was shown during the behavioral learning session. *Temporally unexpected context*: both the two leading images (L_L1_ and L_R1_) and two trailing images (T_L2_ and T_R2_) were expected in terms of spatial context (same as the expected condition), the temporal association was violated (i.e., L_L1_ → T_L2_ and L_R1_ → T_R2_). *Spatially unexpected context*: while the leading image reliably predicted the identity of the trailing image on both the left (L_L1_ → T_L1_) and right (L_R2_ → T_R2_) side independently, thus retaining the expected temporal context, image pairs were not associated in terms of spatial context, neither during the leading images nor during the two trailing images (e.g., L_L1_ and L_R2_ occurring together). *Both unexpected*: shown were four images that do not appeared together in the expected condition. Therefore, the expectation violations occurred in both the temporal and spatial contexts.

To preview our results, we show that spatial and temporal context both modulate sensory processing in key areas of the ventral visual stream, with pronounced reductions in neural responses to stimuli predicted by spatial and temporal context, compared to stimuli occurring in unexpected contexts.

Interestingly, spatial context predictions resulted in a larger suppression than temporal context predictions, suggesting that spatial context may be a more potent modulator of visual processing than temporal context.

## Materials and Methods

### Data and code availability

All data and code used for stimulus presentation and analysis is freely available on the Donders Repository (https://data.donders.ru.nl/login/reviewer-96936509/hUq0EMV2cQaXlzwhl3XLeHsm3q5xbRMZoSX6-YzhpZc).

### Participants

Thirty-three healthy, right-handed participants (13 females, aged 22.36 ± 2.38 years, mean ± SD) were recruited in exchange for monetary compensation (100 Yuan/hour). All participants reported normal or corrected-to-normal vision and were prescreened for MRI compatibility, had no history of epilepsy or cardiac problems.

The experiments reported here were approved by the Institutional Review Board of Psychological Sciences at Hangzhou Normal University and were carried out in accordance with the guidelines expressed in the Declaration of Helsinki. Written informed consent was obtained from all participants. Data from two participants were excluded. Of these two exclusions, one participant’s behavioral performance of the post-scanning task was at chance level, while the other participant showed excessive head motion (i.e., a number of relatively head motion events exceeding 1 mm notably above the group mean).

### Stimuli

The object images were a selection of stimuli from Brady et al. (2008), and also previously used by Richter and de Lange (2019). A subset of 48 full color object stimuli, comprised of 24 electronic objects and 24 non-electronic objects were shown during the present study. For each participant, 24 objects (12 electronics and 12 non-electronics) were pseudo-randomly selected, of which 6 (including 3 electronics) were pseudo-randomly assigned as left leading images, 6 (including 3 electronics) were appointed as right leading images, another 6 (including 3 electronics) served as left trailing images while the remaining 6 (including 3 electronics) acted as right trailing images. Therefore, each specific image could occur in any position or condition (left or right, leading or trailing), thereby minimizing potential biases by specific features of individual object stimuli. Image size was 5° x 5° visual angle presented on a mid-gray background. Stimuli and their association remained the same during the behavioral learning session, MRI scanning and a post-scanning object categorization task. During the behavioral learning session and post-scanning test, object stimuli were presented on an LCD screen (ASUS VG278q, 1920 × 1080 pixel resolution, 60 Hz refresh rate). During MRI scanning, stimuli were displayed on a rear-projection MRI-compatible screen (SAMRTEC SA-9900 projector, 1024 × 768 pixel resolution, 60 Hz refresh rate), visible using an adjustable mirror mounted on the head coil.

### Experimental design

Each participant completed two sessions on two consecutive days. The first session comprised a behavioral learning task while the second session included an fMRI task and a post-scanning object categorization task. While the stimuli and their associations were identical during both sessions, different tasks were employed.

#### Day one - Learning session

Each trial began with a black fixation dot (diameter = 0.4° visual angle) in the center of the screen, participants were asked to maintain fixation on the fixation dot throughout the trial. Two leading images were presented 1.5° visual angle left and right from the central fixation dot for 500 ms, immediately followed by two trailing images, without ISI, at the same locations for 500 ms (**Figure 1a**). Participants were required to count the pairs of same category objects (electronic vs. non-electronic) shown during the leading and trailing images and respond within 2000 ms after trailing image onset by pressing one of three response buttons (corresponding to none, one, or both; see *Pair counting task* below for details). Finally, feedback was presented for 500 ms, followed by a 1000 - 2000 ms ITI. 24 object images (12 electronics and 12 non-electronics) were pseudo-randomly preselected per participant from a pool of images, 12 of which were pseudo-randomly combined into pairs, forming a total of 6 leading image pairs (i.e., the first two images on a trial), while the remaining 6 pairs were used as trailing image pairs (i.e., the second two images on a trial). Crucially, during the learning session, the leading image pair was perfectly predictive of the identity of the trailing image pair [P(trailing pair | leading pair) = 1]. At the same time, the left and right images within both the leading and trailing image pairs were 100% predictive of one another (i.e., pairs always occurred together). Thus resulting in deterministic association in both spatial (co-occurrence) and temporal (sequence) contexts during learning session (see the most left panel in **Figure 1c**). During the learning session each participant performed 5 blocks, with each block comprised of 216 trials, resulting in a total of 180 trials per pair during learning session. The learning session took approximately 60 minutes.

#### Day two – fMRI session

One day after the learning session, participants performed the fMRI session. This session started with one additional block identical to the behavioral learning session, including 216 trials, to renew the learned associations before MRI scanning. During MRI scanning, participants first performed 36 practice trials during acquisition of the anatomical image. The fMRI session was similar to the behavioral learning session, except for the following three modifications. First, a longer ITI of 2000 – 6000 ms (mean = 3000 ms) was used. Second, instead of counting pairs of the same category, participants were required to detect oddball images. Oddballs were the same object images, as shown before, but flipped upside-down, occurring on 10% of trials. Participants were instructed to respond to these target images by pressing a button as quickly as possible, while no response was required during trials without an oddball image. Crucially, whether an image was upside-down was completely randomized and could not be predicted on the basis of the statistical regularities that were present in the image sequences. Third, while the association between images remained the same as during the behavioral learning session, in the fMRI session also unexpected image pairs were shown. In particular, the transition matrices shown in **Figure 1b**, determined how often images were presented together. In 50% of trials, a leading image pair was followed by its expected trailing image pair, identical to the learning session, thus constituting the expected condition. For instance, L_L1_ (leading image, left 1) and L_R1_ (leading image, right 1) served as leading image pair for T_L1_ (trailing image, left 1) and T_R1_ (trailing image, right 1). In the other half of trials, one of the three unexpected conditions (temporally unexpected context, spatially unexpected context, both temporally and spatially unexpected context) occurred with equal possibilities, resulting in 16.67% per unexpected condition. Specifically, for the temporal unexpected context (**Figure 1c** left middle panel), after presenting a leading image pair, one of the other five unmatched trailing image pairs would occur. Thus, while the two images within both the leading and trailing image pair were still expected (i.e., no spatial expectation violation), the temporal sequence of images was unexpected. For example, in this condition L_L1_ and L_R1_ were followed by T_L2_ and T_R2_. For the spatially unexpected context (**Figure 1c** right middle panel), each leading image was followed by its expected trailing image (e.g., L_L1_ → T_L1_ and L_R2_ → T_R2_). However, the two images presented during both the leading and trailing image period were not usually paired; e.g., L_L1_ × L_R2_ and T_L1_ × T_R2_). Thus, in this condition spatial context expectations were violated, while temporal context was expected, thus constituting the spatially unexpected condition. In a final condition, both, spatial and temporal context were violated (**Figure 1c** most right panel). In particular, all four images shown during this condition did not appeared together in the learning session. Crucially, the expectation status only depended on the usual association between the leading image pair and trailing image pair, rather than the frequency or identity of an object image per se. In other words, each object image occurred as expected object and in each unexpected condition. Therefore, all images occurred equally often throughout the experiment, ruling out potential confounds of stimulus frequency or familiarity. Feedback on behavioral performance (accuracy) was provided after each run.

During MRI scanning, each run consisted of 108 trials, including 54 expected trials, 18 temporal context violation trials, 18 spatial context violation trials and 18 trials where both spatial and temporal context were violated. The order of trials was randomized within each run. In total each participant performed 5 runs. Each run lasted ∼12 minutes with 5 null events of 12 s that were evenly distributed across the run, which also served as brief resting periods. The first 8 s of fixation was discarded from analysis. Finally, after MRI scanning, a pair counting task, identical to the learning session was performed outside of the MRI scanner room, which took approximately 20 minutes (see *Pair counting task* below for details).

#### Functional localizer

Following the main task runs during the fMRI session, two functional localizer runs were scanned. These localizer runs were used to define object-selective LOC, and to select voxels that were maximally responsive to the relevant object images. For each participant, the same 12 trailing images that were previously seen in the main task runs and their phase-scrambled version were presented during the localizer. Images were presented at the left and right from the center of screen, corresponding to the location where the stimuli were shown during the main task runs. Each image was shown for 11 s, alternating between the left and right side. Images flashed with a frequency of 2 Hz (300 ms on, 200 ms off). Throughout the localizer, participants were instructed to fixate the fixation dot, while monitoring for an unpredictable dimming of the stimulus (dimming period = 300 ms). Participants responded as quickly as possible by pressing a button. In each run, 4 null events of 11 s were evenly inserted, and each trailing image and its phase-scrambled version was presented two times. The order of trials was fully randomized, except for excluding direct repetitions of the same image. Each participant completed two localizer runs, with each run lasting ∼9.5 minutes. In total each image and its phase-scrambled version was presented 4 times.

#### Pair counting task

Because the oddball detection performed during fMRI scanning does not relate to the underlying statistical regularities, and therefore does not indicate whether statistical regularities were indeed learned, an additional pair counting task was performed after fMRI scanning. In this task, participants were asked to count the number of pairs of the same object category shown on each trial. Participants were further instructed to respond as quickly and accurately as possible. Thus, this task was the same as the task performed during the behavioral learning session, except that the three unexpected conditions were also included. The rationale of this task was to gauge the learning of the object pairs (i.e., statistical regularities) in terms of both temporal and spatial context. Participants could benefit from the knowledge of the associations between the image pairs, as both knowledge about the co-occurrence and temporal sequence would allow for faster responses.

Therefore, the performance difference (e.g., accuracy and reaction time) between the expected condition and each unexpected condition could be considered as an indication for having learnt the underlying statistical regularities. In total, participants performed 360 trials split into 2 blocks, including 180 expected trials, 60 temporally unexpected context trials, 60 spatially unexpected context trials and 60 trials in which both spatial and temporal context were unexpected. The pair counting task took approximately 20 minutes.

### fMRI parameters

Functional and anatomical images were acquired on a 3.0T GE MRI-750 system (GE Medical Systems, Waukesha, WI, USA) at Hangzhou Normal University, using a standard 8-channel headcoil. Functional images were acquired in a sequential (ascending) order using a T2*-weighted gradient-echo EPI pulse sequence (TR/TE = 2000/30 ms, voxel size 2.5 × 2.5 × 2.3 mm, 0.2 mm slice space, 36 transversal slices, 75° flip angle, FOV = 240 mm^2^). Anatomical images were acquired using a T1-weighted inversion prepared 3D spoiled gradient echo sequence (IR-SPGR) (inversion time = 450 ms, TR/TE = 8.2/3.1ms, FOV = 256 × 256 mm^2^, voxel size 1 × 1 × 1 mm, 176 transversal slices, 8° flip angle, parallel acceleration = 2).

## Data analysis

### Behavioral data analysis

Behavioral data from the pair counting task was analyzed in terms of response accuracy and RT. RT was calculated relative to the onset of the trailing image objects. Only trials with correct responses were included in RT analysis. Additionally, we excluded trials with RTs shorter than 200 ms (0.82%) or more than three standard deviations above the subject’s mean response time (0.49%). RT and accuracy data for expected and unexpected trailing image trials were averaged separately per participant and across subjects subjected to a paired t test. The effect size was calculated in terms of Cohen’s *d*_z_ for all paired t-test, while partial eta-squared (*η*^2^) was used for indicating effect sizes in the repeated measures ANOVA (Lakens, 2013).

### fMRI data preprocessing

fMRI data preprocessing was performed using FSL 6.0.1 (FMRIB Software Library; Oxford, UK; www.fmrib.ox.ac.uk/fsl; Smith et al., 2004, RRID:SCR_002823). The preprocessing pipeline included brain extraction (BET), motion correction (MCFLIRT), slice timing correction (Regular up), temporal high-pass filtering (128 s), and spatial smoothing for univariate analyses (Gaussian kernel with FWHM of 5 mm). Functional images were registered to the anatomical image using FSL FLIRT (BBR) and to the MNI152 T1 2 mm template brain (linear registration with 12 degrees of freedom). Registration to the MNI152 template brain was only applied for whole-brain analyses, while all ROI analyses were performed in each participant’s native space in order to minimize data interpolation.

### Whole brain analysis

To estimate the BOLD response to expected and unexpected stimuli across the entire brain, FSL FEAT was used to fit voxel-wise general linear models (GLM) to each participant’s run data in an event-related approach. In the first level GLMs, expected and three unexpected image object trials were modeled as four separate regressors with a duration of one second (the combined duration of leading and trailing image pairs), and convolved with a double gamma hemodynamic response function. An additional nuisance regressor for oddball trials (upside-down images) was added. Additionally, first-order temporal derivatives for the five regressors, and 24 motion regressors (FSL’s standard + extended motion parameters) were also added to the GLM. To quantify the main effects of spatial and temporal expectation suppression, we contrasted unexpected regressors and the expected regressors for spatial and temporal context separately (i.e., temporal context expectation suppression = BOLD_Temporal unexpected_ + BOLD_Both unexpected_ - BOLD_Spatial unexpected_ - BOLD_Both expected_; spatial context expectation suppression = BOLD_Spatial unexpected_ + BOLD_Both unexpected_ - BOLD_Temporal unexpected_ - BOLD_Both expected_). Data were combined across runs using FSL’s fixed effect analysis. For the across-participants whole-brain analysis, FSL’s mixed effect model (FLAME 1) was used. Multiple-comparison correction was performed using Gaussian random-field based cluster thresholding.

The significance level was set at a cluster-forming threshold of z > 3.1 (i.e., *p* < 0.001, two-sided) and a cluster significance threshold of *p* < 0.05.

### Regions of interest (ROIs) analysis

ROI analyses were conducted in each participant’s native space. Primary visual cortex (V1), object-selective lateral occipital complex (LOC), and temporal occipital fusiform cortex (TOFC) were chosen as the three ROIs (see *ROI definition* below) for analysis, based on two previous studies that used a similar experimental design (Richter et al., 2018; Richter and de Lange, 2019). The mean parameter estimates were extracted from each ROI for the expected and unexpected conditions separately. For each ROI, these data were submitted to a two-way repeated measures ANOVA with temporal context (expected vs. unexpected) and spatial context (expected vs. unexpected) as factors.

### ROI definition

All ROIs were defined using independent data from the localizer runs. Specifically, V1 was defined based on each participant’s anatomical image, using Freesurfer 6.0 to define the gray–white matter boundary and perform cortical surface reconstruction (recon-all; Dale et al., 1999; RRID:SCR_001847). The resulting surface-based ROI of V1 was then transformed into the participant’s native space and merged into one bilateral mask. Object selective LOC was defined as bilateral clusters, within anatomical LOC, showing a significant preference for intact compared to scrambled object stimuli during the localizer run (Kourtzi and Kanwisher, 2001; Haushofer et al., 2008). To achieve this, intact objects and scrambled objects were modeled as two separate regressors in each participant’s localizer data. The temporal derivatives of all regressors and the 24 motion regressors were also added to fit the data. Finally, the contrast of interest, objects minus scrambles, was constrained to anatomical LOC. In order to create the TOFC ROI mask, the anatomical temporal-occipital fusiform cortex mask from the Harvard-Oxford cortical atlas (RRID:SCR_001476), distributed with FSL, was further constrained to voxels showing a significant conjunction inference of expectation suppression on the group level in Richter et al. (2018) and Richter and de Lange (2019). The resulting mask was then transformed from MNI space to each participant’s native space using FSL FLIRT. Finally, the 200 most active voxels in each of the three ROI masks were selected for further statistical analyses. To this end, the contrast interest between the left and right hemisphere in V1 (including both the intact and scrambled images) was calculated, while in LOC and TOFC, the contrast interest between the intact images and the scrambled images was calculated based on the localizer data. The resulting z-map of this contrast was then averaged across runs. Finally, we selected the 200 most responsive voxel from this contrast. In order to verify that our results did not depend on the a priori defined, but arbitrary number of voxels in the ROI masks, we repeated all ROI analyses with masks ranging from 50 to 500 voxels in steps of 50 voxels.

### Bayesian analysis

In order to further evaluate any non-significant results, and arbitrate between an absence of evidence and evidence for the absence of an effect, the Bayesian equivalents of the above outlined analyses were additionally performed. JASP 0.10.2 (JASP Team, 2019, RRID:SCR_015823) was used to perform all Bayesian analyses, using default settings. Thus, for Bayesian t-tests a Cauchy prior width of 0.707 was chosen. Qualitative interpretations of Bayes Factors are based on criteria by Lee and Wagenmakers (2014).

## Results

We exposed participants to statistical regularities by presenting two successive object image pairs in which the leading image pairs predicted the identity of the trailing image pairs. The identities of the image pairs were also predictable in terms of their spatial context; i.e., simultaneously shown left and right images occurred together. Subsequently, in the MRI scanner, participants were shown the same predictable object image pairs (expected condition), but additional expectation violations were introduced. In particular, either the temporal context was violated, the spatial context was violated, or both contexts were violated (see **Figure 1c**).

### Stronger modulation of spatial context than temporal context on sensory processing throughout the ventral visual stream

In order to assess the consequences of violating temporal and spatial context expectations we performed a two-way repeated measures ANOVA with temporal context (expected vs. unexpected) and spatial context (expected vs. unexpected) as factors, within our a prior defined ROIs: primary visual cortex (V1), object-selective lateral occipital complex (LOC), and temporal occipital fusiform cortex (TOFC). In higher visual areas, LOC and TOFC, we observed a significant decrease in BOLD responses when stimuli were expected in terms of their spatial context (**Figure 2a**; LOC: *F*_(1, 32)_ = 31.389, *p* = 3.0e-6, *η*^2^= 0.495; TOFC: *F*_(1, 32)_ = 23.083, *p* = 3.5e-5, *η*^2^ = 0.419). In other words, when two stimuli frequently co-occurred, thus making them expected in this pair, they elicited reduced sensory responses in ventral visual areas. Furthermore, we found a similar suppression of neural responses by temporal context expectations in TOFC (*F*_(1, 32)_ = 10.805, *p* = 0.0025, *η*^2^= 0.252), but not in LOC (*F*_(1, 32)_ = 1.266, *p* = 0.2689, *η*^2^= 0.038). That is, in TOFC, if a pair of stimuli was expected given the preceding stimulus pair, the elicited BOLD response was suppressed compared to the response to the same pair occurring in an unexpected temporal sequence. No interaction between temporal and spatial context was found in either LOC or TOFC (LOC: *F*_(1, 32)_ = 0.111, *p* = 0.7412, *η*^2^= 0.003; TOFC: *F*_(1, 32)_ = 0.064, *p* = 0.8013, *η*^2^= 0.002). Thus, the suppression of neural responses induced by temporal expectations was not modulated by spatial context expectations, and vice versa.

**Figure 2.**
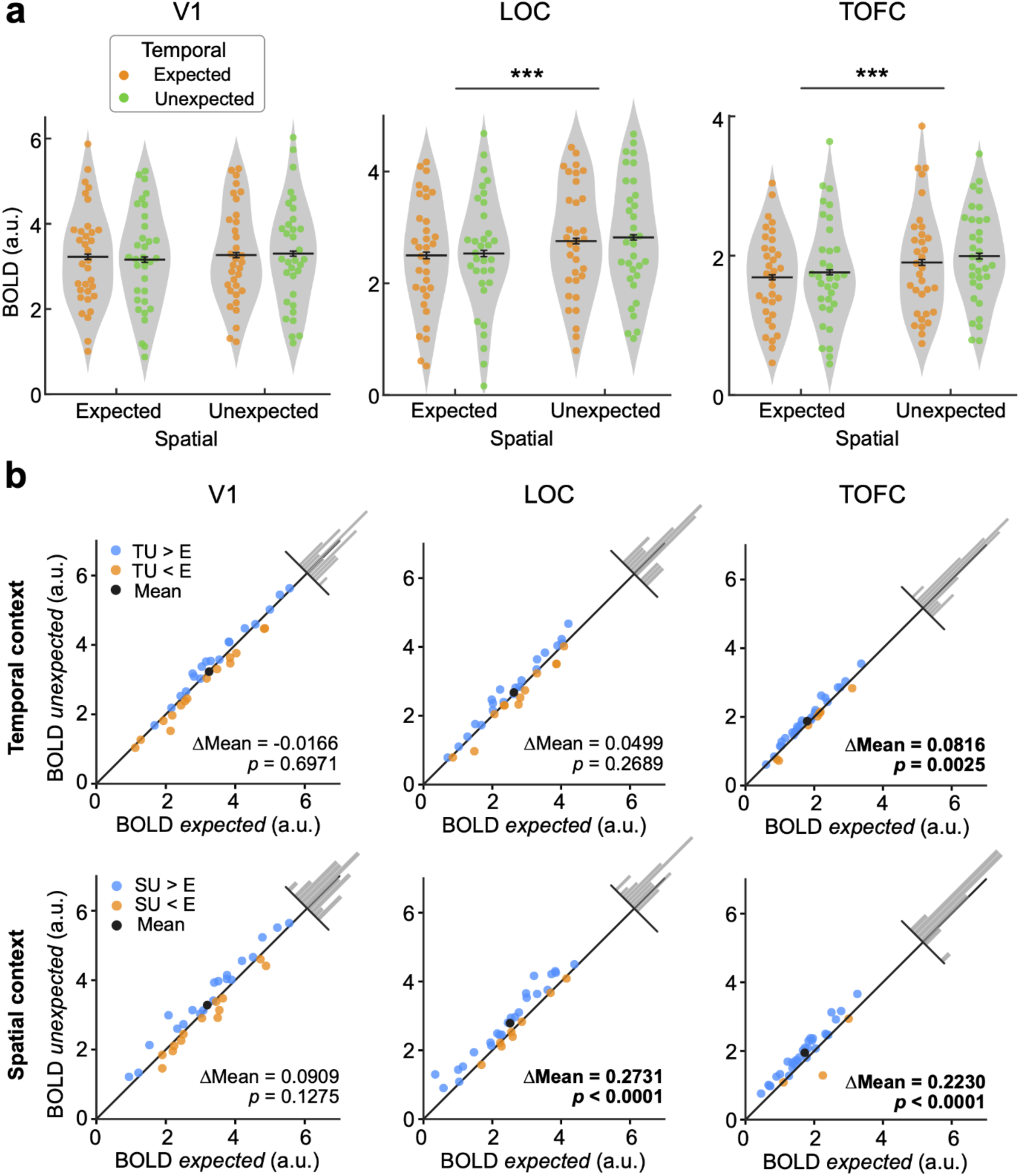
Expectation suppression within V1, LOC and TOFC. **(a)** Parameter estimates for responses to expected and unexpected images pairs. In both LOC and TOFC, BOLD responses to spatially expected image pairs were significantly attenuated compared to unexpected image pairs. Furthermore, a reliable suppression of responses by temporal context expectations was observed in TOFC. No modulation of BOLD responses by expectations was found in V1. Each dot denotes an individual participant and the black line is the mean across participants. Error bars denote ±1 within-subject SEM. \**p* < 0.05, \*\**p* < 0.01, \*\*\**p* < 0.001. **(b)** BOLD responses evoked by unexpected and expected context within V1 (left column), LOC (middle column) and TOFC (right column). The upper row represents the BOLD contrast between the temporally unexpected context and expected context, averaged across the spatially expected and unexpected context. The bottom row represents the BOLD contrast between the spatially unexpected context and expected context, averaged across the temporally expected and unexpected context. Blue and yellow dots represent individual participants. Blue indicates expectation suppression (unexpected > expected), yellow indicates expectation enhancement (unexpected < expected), and black indicates the mean of all subjects. ΔMean is equal to the difference of BOLD response between the unexpected and expected condition. The inset histogram shows the distribution of deviations from the unity line.

In a post-hoc analysis we compared the magnitude of neural suppression induced by temporal and spatial context predictions. In LOC and TOFC spatial context expectations resulted in a larger suppression than temporal expectations (LOC: *t*_(32)_ = 2.870, *p* = 0.0072, Cohen’s *d*_z_ = 0.835; TOFC: *t*_(32)_ = 2.575, *p* = 0.0149, Cohen’s *d*_z_ = 0.691), thus suggesting that spatial context may be a stronger modulator of visual responses than temporal context.

Perhaps surprisingly, we did not find any reliable modulation of neural responses by temporal or spatial context predictions in V1 (spatial context: *F*_(1, 32)_ = 2.448, *p* = 0.1275, *η*^2^= 0.071; temporal context: *F*_(1, 32)_ = 0.154, *p* = 0.6971, *η*^2^= 0.005; spatial context by temporal context interaction: *F*_(1, 32)_ = 0.627, *p* = 0.4342, *η*^2^ = 0.019). Indeed, in V1, Bayesian analyses yielded moderate evidence for the absence of a modulation of neural responses by temporal context violations (temporally unexpected context vs. expected context: BF_10_ = 0.141), and anecdotal support for the absent of an effect when spatial context was violated (spatially unexpected context vs. expected context: BF_10_ = 0.388). Thus, in V1 expectations, in terms of temporal or spatial context, did not appear to modulate sensory responses. In contrast, in higher visual areas a suppression of responses to expected stimuli was observed both for temporal and spatial contexts.

To ensure that our results were not dependent on the a prior but arbitrarily chosen mask sizes of the ROIs, we repeated the analyses for ROIs of sizes ranging from 50 to 500 voxels in step of 50 voxels. Results were qualitatively identical to those mentioned above (**Figure 2a**) for all ROI sizes within all three ROIs (V1, LOC, TOFC), indicating that our results do not depend on ROI size, but well represent results within the ROIs.

A complementary whole-brain analysis was performed to investigate the effect of temporal context and spatial context outside of our predefined ROIs. Results are illustrated in **Figure 3**. In accordance with our ROI analysis, spatial expectations were associated with significantly suppressed neural responses throughout the ventral visual stream. Additional clusters of expectation suppression were evident outside the ventral visual stream, including bilateral frontal gyrus, bilateral precentral gyrus, bilateral frontal operculum and insular cortex, as well as the paracingulate gyrus. In contrast, no reliable modulation by temporal context expectation was found outside of our predefined ROIs in the whole-brain analysis. Thus, temporal context expectations were only evident in the ROI analysis, but too small or hidden by interindividual variability to be detected in the whole-brain analysis (note: ROI masks were individually defined for each participant; also see Materials and Methods, ROI definition).

**Figure 3.**
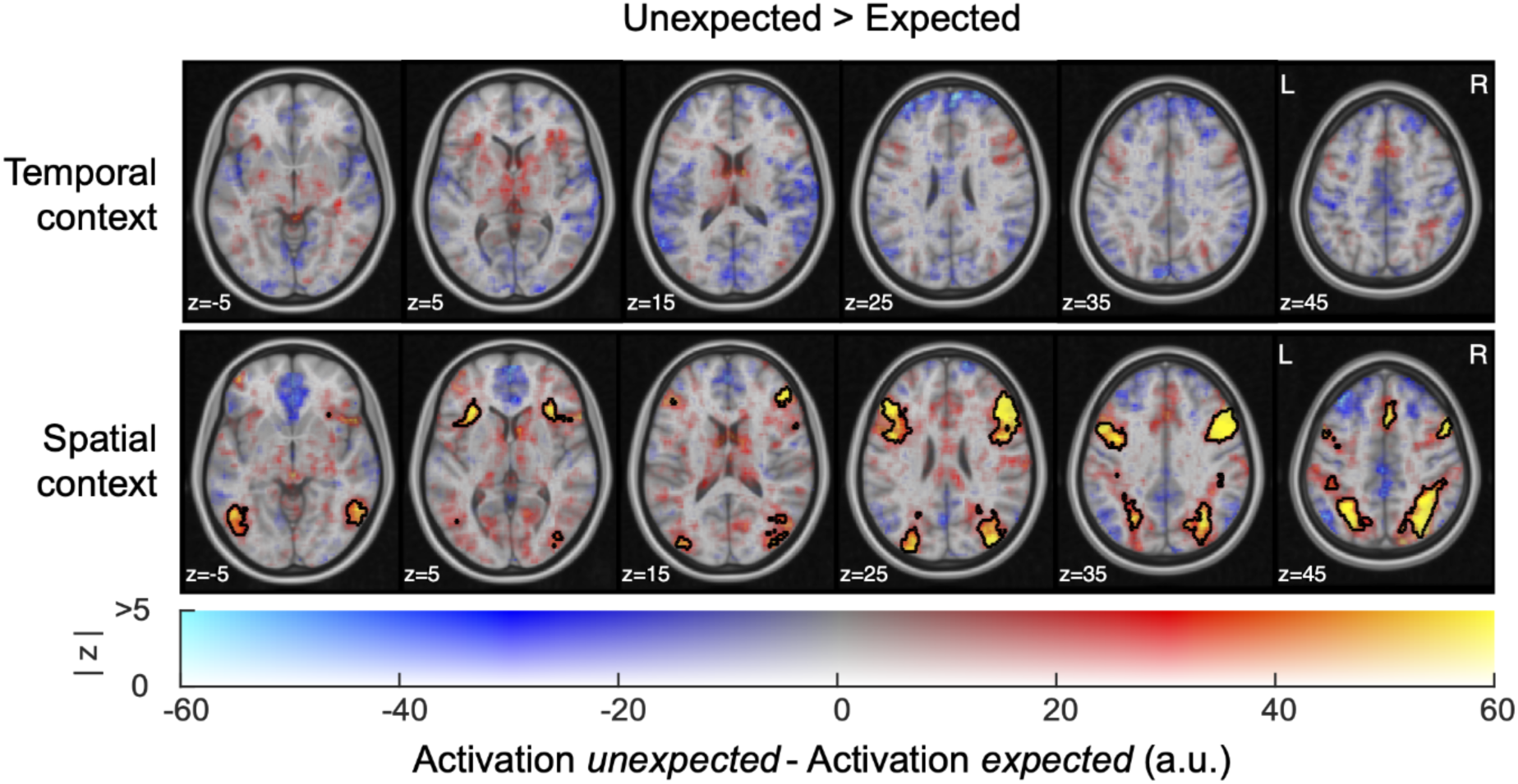
Expectation suppression across cortex for temporal and spatial contexts. Displayed are parameter estimates for unexpected minus expected image pairs overlaid onto the MNI152 2 mm anatomical template. Color represents the unthresholded parameter estimates: red-yellow clusters denote expectation suppression, blue-cyan clusters indicate expectation enhancement; opacity indicates the z statistics of the contrast. Black contours outline statistically significant clusters (Gaussian random field cluster corrected). No significant clusters were found for the main effect of temporal context (upper row). The main effect of spatial expectation (bottom row) shows significant clusters of expectation suppression in parts of the ventral visual stream (LOC, TOFC), as well as bilateral frontal gyrus, bilateral precentral gyrus, bilateral frontal operculum and insular cortex, and paracingulate gyrus.

### Expectations facilitate object categorization

In addition to the neural effects of expectations, we also examined whether expectations facilitated behavioral responses. During a post-scanning object categorization task, participants were asked to count the number of object pairs of the same category shown as leading and trailing image pairs (i.e., 0, 1 or 2 pairs could be of the same category). In order to fulfill this task, as quickly and accurately as possible, participants could benefit from the knowledge of the underlying statistical regularities – both in terms of co-occurrence (spatial) and sequence (temporal) prediction. In line with our hypothesis, RTs and accuracy of responses (**Figure 4**) were affected by expectations, in both temporal (RT: *t*_(32)_ = 4.891, *p* = 6.9e-6, Cohen’s *d*_z_ = 0.851; accuracy: *t*_(32)_ = 4.924, *p* = 6.1e-6, Cohen’s *d*_z_ = 0.857) and spatial contexts (RT: *t*_(32)_ = 11.670, *p* = 1.3e-17, Cohen’s *d*_z_ = 2.031; accuracy: *t*_(32)_ = 10.224, *p* = 3.7e-15, Cohen’s *d*_z_ = 1.780). Thus, participants learned and benefitted from both spatial and temporal context predictions.

**Figure 4.**
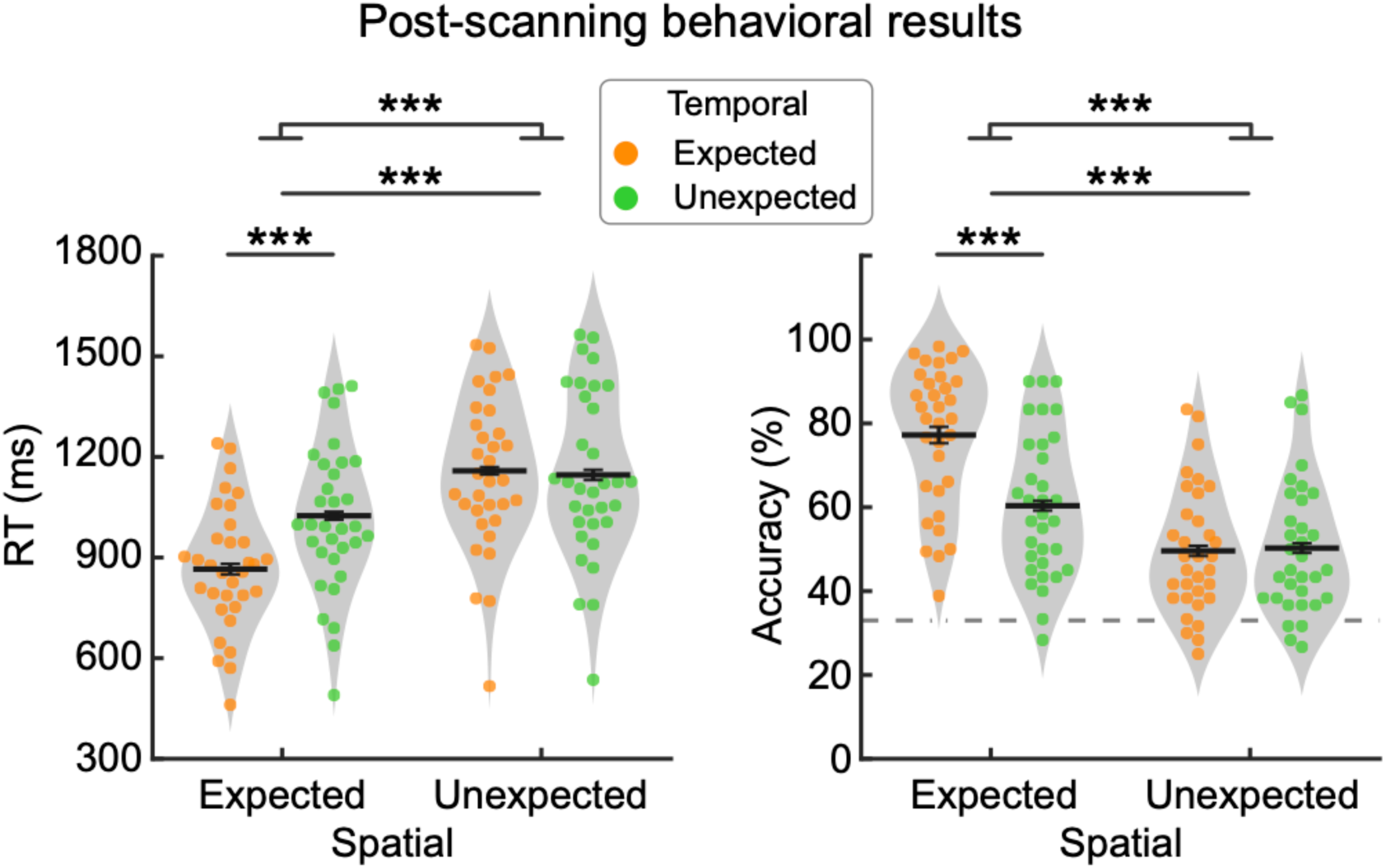
Behavioral data indicate statistical learning. Reaction time (left) and accuracy (right) are plotted for expected and unexpected conditions in temporal (dot color) and spatial contexts (abscissa), respectively. Behavioral responses in the spatially expected condition are significantly faster and more accurate than in the unexpected condition. Temporally expected stimulus pairs also result in faster and more accurate responses, however this effect is only present when spatial expectations were met. Dashed horizontal gray line indicates chance level accuracy (33.33%). Dots represent single subject data. Black line is the mean across participants. Error bars denote ±1 within-subject SEM. \*\*\**p* < 0.001.

Interestingly, participants were faster and more accurate in response to objects predicted by the temporal sequence only when the spatial context was expected as well (RT: *t*_(32)_ = 9.329, *p* = 1.2e-10, Cohen’s *d*_z_ = 1.624; accuracy: *t*_(32)_ = 7.649, *p* = 1.0e-8, Cohen’s *d*_z_ = 1.332), but not when the spatial context was unexpected (RT: *t*_(32)_ = 0.269, *p* = 0.7898, Cohen’s *d*_z_ = 0.047, BF_10_ = 0.193; accuracy: *t*_(32)_ = 0.566, *p* = 0.5755, Cohen’s *d*_z_ = 0.099, BF_10_ = 0.216). The robustness of this distinct pattern of facilitation effect was statistically confirmed by an interaction analysis (RT: *F*_(1, 32)_ = 38.787, *p* = 5.6e-7, *η*^2^= 0.548; accuracy: *F*_(1, 32)_ = 46.337, *p* = 1.1e-7, *η*^2^= 0.592). Moreover, when a stimulus was expected by spatial context, participants showed faster and more accurate responses, irrespective of whether the temporal context was expected (RT: *t*_(32)_ = 13.977, *p* = 3.6e-15, Cohen’s *d*_z_ = 2.433; accuracy: *t*_(32)_ = 10.883, *p* = 2.7e-12, Cohen’s *d*_z_ = 1.894) or unexpected (RT: *t*_(32)_ = 5.838, *p* = 1.7e-6, Cohen’s *d*_z_ = 1.016; accuracy: *t*_(32)_ = 6.279, *p* = 4.9e-7, Cohen’s *d*_z_ = 1.093).

In sum, behavioral performance was reliably facilitated by spatial context, resulting in faster and more accurate responses. On the other hand, expected temporal sequences also aided in faster and more accurate responses, however only when the spatial context was expected. These results may suggest that participants grouped pairs of objects, and predicted the upcoming pair of objects, instead of individual sequences of objects on the left and right side separately.

### Spatial and temporal context expectations modulate neural responses in similar cortical areas

Given the modulation of neural responses by temporal context in TOFC in our ROI analysis (**Figure 2a**), and the reliablity of expectation suppression reported in previous studies investigating temporal context violations (Turk-Browne et al., 2009; Meyer and Olson, 2011; Richter and de Lange, 2019), it is perhaps surprising that we did not find evidence of temporal expectation effects in the whole-brain analysis (**Figure 3**). Potential explanations why temporal context violations did show little effect in the present study will be discussed in more detail later (see Discussion).

However, in order to further compare spatial and temporal predictions, it could be informative to compare the localization of the here reported spatial expectation suppression with temporal expectation suppression shown in previous studies. In a conjunction analysis, we investigated the overlap of expectation suppression between previously reported temporal expectation suppression from Richter and de Lange (2019) and the present spatial expectation violation. Results illustrated in **Figure 5**, show clusters of overlapping expectation suppression between temporal and spatial context expectations throughout parts of the ventral visual stream, and several non-sensory areas, including middle and inferior frontal gyrus, precentral gyrus. Thus, spatial context expectations, as observed here, and temporal context expectations, as reported by Richter and de Lange (2019), are evident in a similar neural network, thereby suggesting that a comparable neural mechanism may underlie both spatial and temporal context predictions.

**Figure 5.**
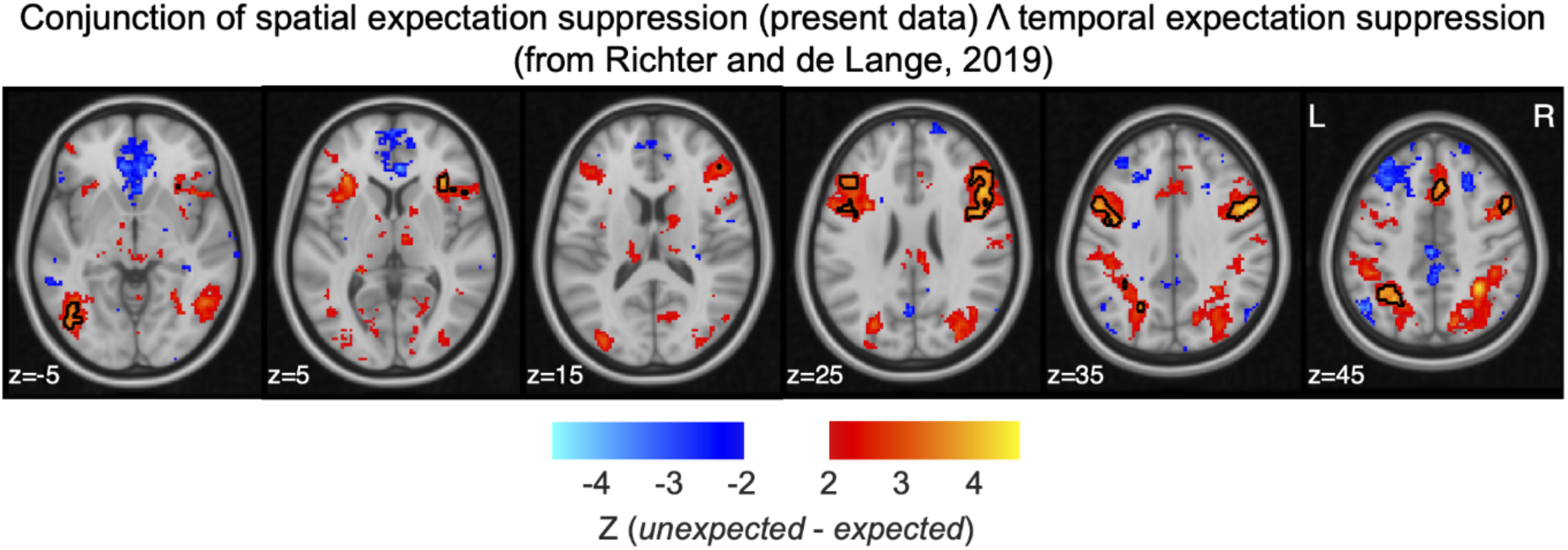
Displayed are z statistics of the contrast between unexpected and expected of a conjunction inference between data from the spatial context violation and data from a temporal context violation effect from Richter and de Lange (2019). Red-yellow clusters denote expectation suppression. Significant overlaps in the localization of expectation suppression include clusters in parts of the ventral visual stream, middle and inferior frontal gyrus, and precentral gyrus.

## Discussion

Both spatial and temporal context play an important role in visual perception and behavior (Schwartz et al., 2007). The present study investigated the neural consequences of violations of expectations derived from spatial and temporal context, across the ventral visual stream. To this end, we exposed participants to two forms of statistical regularities, making stimuli predictable in terms of spatial context (co-occurrence of stimuli at specific locations) and temporal context (specific temporal sequence of stimuli). While we measured brain activity to these stimuli, image transitions were not task relevant, and thus any neural modulations by spatial and temporal context were not dependent on task-relevance of the underlying statistical regularities. We found a reliable and wide-spread activity modulation in the ventral visual stream, including LOC and TOFC, as a function of spatial context. In particular, when stimuli frequently co-occurred neural responses were suppressed compared to the response to the same stimulus co-occurring with another stimulus, even though all stimuli were equally familiar and always occurred at the same spatial location. Temporal context (i.e., predictability of stimulus sequence) also modulated neural responses in TOFC, again evident as a suppression of responses to expected stimuli. Interestingly, while the two forms of context modulated overlapping regions, the activity modulation by spatial context was much stronger and more wide-spread than the modulation by temporal context. Thereby our results extend previous studies (e.g., Summerfield et al., 2008; Alink et al., 2010; Kok et al., 2012; Richter et al., 2018; Richter and de Lange, 2019) by demonstrating that spatial and temporal context priors may modulate neural responses in a similar fashion and within the same cortical network. However, at least in the visual system spatial context appears to be a more potent modulator of perceptual processing than temporal context.

### Spatial and temporal context facilitate behavior

Our data showed a substantial and robust facilitation of behavioral responses by both spatial and temporal contexts. During a post-scanning test, requiring participants to count stimulus pairs of the same category (i.e., both electronic, or both non-electronic stimuli), spatial and temporal context strongly modulated behavioral performance (**Figure 4**). Specifically, responses were faster and more accurately to stimuli presented in a spatially and temporally expected context, and the violation of either context increased RTs and decreased response accuracy – with larger decrements for spatial context violations. Crucially, the benefit of temporally expected contexts was only observed when the spatial context was expected. However, performance enhanced by spatially expected contexts was evident irrespective of whether temporal context expectations were confirmed or violated.

Thus, our data show that participants can in principle learn and benefit from both spatial and temporal statistical regularities. However, our results also suggest that our participants may have grouped simultaneously presented objects into image pairs, which combined predicted the next image pair. That is, even though object stimuli on the left and right side predicted the identity of the next stimulus independently, even when spatial configuration were unexpected, these statistical regularities may not have been learned, or the resulting predictions may not have been instantiated. These results may suggest a preference for spatial over temporal grouping in vision. However, it is important to note here that a strategy of grouping spatial pairs may have partially been induced by the same-different category counting task during learning, which specifically requires participants to make a judgment about the groups of objects.

### Spatial and temporal context modulate sensory processing in the ventral visual stream

Our fMRI results show that sensory responses in object selective visual areas (LOC and TOFC) are suppressed, if stimuli occur in expected spatial contexts compared to unexpected spatial contexts. In other words, stimuli that frequently co-occur evoked reduced sensory responses relative to the same stimuli presented in less frequently co-occurring configurations. Note, that the frequency of the individual stimuli occurring were equal, thereby excluding potentially confounding effects of stimulus frequency or familiarity. Moreover, during MRI scanning predictions were task-irrelevant, thus suggesting that predictions were formed and modulated neural responses automatically.

The suppression of neural responses by spatial predictions matches key characteristics of expectation suppression, a phenomenon previously described in terms of suppressed sensory responses to stimuli expected by virtue of their temporal context; i.e., a leading image predicting the identity of a trailing image (den Ouden et al., 2009; Meyer and Olson, 2011; Richter et al., 2018; Richter and de Lange, 2019). In line with previous studies, we also found a suppression of sensory responses by temporal context in TOFC. That is, stimuli in expected temporal sequences elicited suppressed BOLD responses compared to stimuli in unexpected temporal sequences.

Moreover, using a conjunction analysis we showed that the here observed spatial context suppression is evident in similar cortical areas as previously reported suppression by temporal context expectations (e.g., den Ouden et al., 2009; Turk-Browne et al., 2009, 2010; Gheysen et al., 2011; Meyer and Olson, 2011; Richter et al., 2018; Richter and de Lange, 2019). Interestingly, this overlap in cortical regions was not limited to object selective visual cortex, but also included several non-sensory areas, such as inferior frontal gyrus. Combined these results suggest that spatial and temporal contexts can have similar modulatory effects on neural processing, thereby implying that the neural mechanism underlying contextual prediction effects may be independent of the type of prediction – temporal or spatial contexts. In agreement with this suggestion, Karuza et al. (2017) reported similar neural modulations, and comparable correlations of these modulations with behavior, during learning of spatial regularities as previously reported for statistical learning of temporal (sequence) regularities (e.g., Turk-Browne et al., 2009, 2010; Gheysen et al., 2010, 2011; Schapiro et al., 2014). Thus, the available data suggest that the neural architecture and computations underlying different types of context predictions may largely overlap, evident in similar modulations of both behavioral and neural responses.

### Stronger modulations of neural responses by spatial context than temporal context

While the present data showed a joint modulation of neural responses by spatial and temporal context, the modulation by temporal context was relatively modest and significantly smaller than the modulation by spatial context. Initially, these results may be surprising given the multitude of previous studies reporting strong and extensive modulations of sensory responses by temporal context predictions across the ventral visual stream (Turk-Browne et al., 2009, 2010; Gheysen et al., 2010; Meyer and Olson, 2011; Tobia et al., 2012a, 2012b; Tremblay et al., 2013; Plante et al., 2015; Richter et al., 2018; Richter and de Lange, 2019).

These previous studies however lacked spatial context, presenting single stimuli in isolation.

Vision is particularly apt to handle simultaneous inputs and the spatial structure between these stimuli (Saffran, 2002). Audition on the other hand shows a remarkable sensitivity to the temporal structure of inputs (Kubovy, 1988; Conway and Christiansen, 2009). Indeed, such modality specific constraints can affect the manner in which stimuli are processed (Mahar et al., 1994; Repp and Penel, 2002), maintained in working memory (Penney, 1989; Collier and Logan, 2000) and learned (Handel and Buffardi, 1969; Saffran, 2002; Conway and Christiansen, 2009). Thus, modality specific biases in the visual system may result in an emphasis on spatial configurations and hence a stronger modulation of neural responses by spatial than temporal context predictions.

Our behavioral results also support the notion that spatial predictions were more readily acquired and utilized than temporal predictions. In particular, only when spatial configurations were expected temporal predictions facilitated behavioral responses. Thus, in the present data, and possibly vision in general, spatial regularities appear to take precedence over temporal statistical regularities, resulting in a larger magnitude of behavioral and neural modulations by spatial compared to temporal context.

### No modulation of neural responses by prediction in primary visual cortex

Surprisingly, we found no modulation by predictions in V1, unlike in some previous studies (e.g., Kok et al., 2012; Richter and de Lange, 2019). It is possible that, because expectations constitute a top-down modulation, likely originating from beyond visual cortex (Hindy et al., 2019), its effect might be less pronounced in V1 compared to higher visual areas. Indeed, in previous studies prediction effects appear to reduce in magnitude in lower visual areas (e.g. see Figure 1A in Richter and de Lange, 2019). Moreover, it is possible that spatial arrangements of object stimuli were too complex to yield specific predictions relevant to the response properties of neural assemblies in V1. That is, predictions in our study constitute arrangements and sequences of full color object images, thus particularly depending on object selective cortical areas. Hence, arrangements of stimuli exploiting the neural tuning in V1, such pairs of oriented grating stimuli may result in prediction induced modulations in V1. Thus, the absence of expectation suppression in V1 observed here may be a consequence of the utilized stimuli and experimental design.

## Conclusion

In conclusion, our data suggest that temporal and spatial statistical regularities jointly facilitate behavioral responses, leading to faster and more accurate responses. At the same time, predictions based on both forms of contexts modulate sensory responses, resulting in a suppression of responses to expected stimuli in a similar cortical network, including object selective visual cortex. However, spatial context appears a more potent modulator within the visual system, resulting in larger modulations of neural responses by spatial compared to temporal context.

## Author contributions

T.H., D.R., Z.W., and F.P.d.L. designed research; T.H. performed research; T.H. and D.R. analyzed data; T.H. and D.R. wrote the first draft of the paper; T.H., D.R., Z.W., and F.P.d.L. edited the paper.

## Acknowledgements

This work was supported by the National Natural Science Foundation of China Grant 31371133 to Z.W., The Netherlands Organisation for Scientific Research Vidi Grant 452-13-016 to F.P.d.L., the EC Horizon 2020 Program ERC Starting Grant 678286 “Contextvision” to F.P.d.L, the James S.McDonnell Foundation 220020373 to F.P.d.L and the China Scholarship Council (CSC; 201608330264) to T.H.. We thank Wanlu Fu and Zehao Huang for assistance with data collection.

